# Complete Genome Sequence of Six *Pseudomonas aeruginosa* Bacteriophages aim for research and treatments

**DOI:** 10.1101/2024.09.05.611452

**Authors:** Amit Rimon, Ortal Yerushalmy, Jonathan Belin, Sivan Alkalay-Oren, Shunit Coppenhagen-Glazer, Ronen Hazan

## Abstract

**Introduction:** *Pseudomonas aeruginosa* is a major opportunistic pathogen associated with healthcare-associated infections. The rise of antibiotic-resistant strains necessitates alternative treatment strategies, with bacteriophage therapy being a promising approach.

**Methods:** Six bacteriophages were isolated from sewage samples. Phage isolation involved centrifugation, filtration, and plaque assays. The morphology of each was examined using Transmission Electron Microscopy (TEM). Genomic DNA was sequenced analyzed and compared. Phages lytic activity was assessed using growth curve analysis.

**Results:** The six phages displayed distinct genomic and morphological characteristics, in three genomic clusters. No known virulence or antibiotic resistance genes were detected, indicating their safety for therapeutic use. TEM analysis revealed diverse morphologies, with some phages belonging to the Siphoviridae family and others to the Myoviridae family. Lysogenic phages demonstrated less effective lytic activity.

**Conclusion:** Some of these phages are promising candidates for the research of phage therapy efficacy, and the lytic phages can be used against *P. aeruginosa* infections.

## Introduction

*Pseudomonas aeruginosa* is a notorious opportunistic pathogen associated with healthcare-associated infections, particularly in immunocompromised patient^1^. Its ability to develop resistance to multiple antibiotics poses a significant challenge in clinical settings, leading to a pressing need for alternative treatment strategies. The rise of antibiotic-resistant strains necessitates alternative treatment strategies, with bacteriophage therapy being a promising approach^2,3^.

Bacteriophages, or phages, are viruses that specifically infect and kill bacteria. They offer a targeted method to combat bacterial infections, which is especially valuable in the era of increasing antibiotic resistance^4^. Recent studies have shown that phage therapy can be an effective means of controlling *P. aeruginosa* infections, particularly in cases where traditional antibiotics fail in humans^3,5,6^ and in animals^7^.

In the IPTC, we have a collection of approximately 70 anti-*P. aeruginosa* phages ^8^. This study focuses on six newly isolated phages from this collection: PAShipCat1, PACT201, PADD, PAB1, PAB2, and PAKlein3. These phages are currently being evaluated for their efficacy in combating *P. aeruginosa* infections.

In this study, we present and discuss the genomic and functional characteristics of these phages. Our research aims to provide a detailed characterization that could pave the way for their potential use in therapeutic applications, offering an alternative to traditional antibiotics in the fight against resistant strains of *P. aeruginosa*.

## Methods

### Phage Isolation and Purification

The phages were isolated from sewage samples collected from the decontamination facility near West Jerusalem as part of the Israeli Phage Bank collection. PAShipCat1, PADD, PAB1, and PAB2 were isolated using *Pseudomonas aeruginosa* PA14^9^ as the host. PAKlein3 was isolated using *Pseudomonas aeruginosa* PA01,^9^ and PACT201 was isolated using the clinical isolate strain PAw3151237 (Table 1).

**Table 1.** Characteristics of Phage Genomes. This table details various genomic characteristics of different phages, including the phage name, bacterial host, genome length, GC content (%), number of coding sequences (CDS), potential for lysogenicity determined by the presence of repressor proteins, and the presence of phage genomes in bacterial genomes identified through BLAST comparisons. Potential for genome circularity as determined as described in the methods section. ^1^Determined by performing BLAST analysis, identifying the closest matches based on query coverage and sequence identity ^2^*Viruses; Duplodnaviria; Heunggongvirae; Uroviricota;* ^3^These genomes have the potential to be circular.

| Phage Name | *Pseudomonas aeruginosa* strain | Genome size | % GC | # CDS | Potential lysogenicity | Topology | GenBank accession number | Most proximal phage<sup>1</sup> |  |  |  |  |  |
| --- | --- | --- | --- | --- | --- | --- | --- | --- | --- | --- | --- | --- | --- |
|  |  |  |  |  |  |  |  | Phage name | % Query cover | % Identity | E value | Taxonomy<sup>2</sup> | GenBank accession number |
| PAShipCat1 | PA14<sup>27</sup> | 50,602 | 59.3 | 82 | Yes | Circular<sup>3</sup> | PP067092 | vB_Pae_SG_WM_Sew_P27 | 61 | 98.52 | 0 | Caudoviricetes | PP740689 |
| PADD |  | 67,298 | 55.7 | 70 | No | Circular<sup>3</sup> | PP931176 | 6917 | 98 | 98.21 | 0 | Caudoviricetes; Pbnavirus | OL362268 |
| PAB1 |  | 51,201 | 55.6 | 63 | No | Linear | PP931177 | bmx-p2 | 99 | 97.92 | 0 | Caudoviricetes; Pbnavirus | OQ319932 |
| PAB2 |  | 45,045 | 58.4 | 54 | No | Linear | PP931178 | Quinobequin-P09 | 95 | 93.22 | 0 | Caudoviricetes; Queuovirinae; Nipunavirus; | NC_072501 |
| PAKlein3 | PA01<sup>27</sup> | 56,901 | 63.4 | 71 | No | Linear | PP931173 | PaeS_PA01_Ab18 | 98 | 96.17 | 0 | Caudoviricetes; Mesyanzhinovviridae; Bradleyvirinae; Abidjanvirus | NC_026594 |
| PACT201 | PAw3151237 | 57,971 | 59.1 | 64 | Yes | Circular<sup>3</sup> | PP931175 | vB_PaeS-D14Q | 39 | 90.81 | 0 | Caudoviricetes | BK061590 |

Briefly, the samples were dissolved in LB broth and mixed with overnight-grown *P. aeruginosa* at 37°C. The culture was then centrifuged (7,100 × g, 10□min), and the supernatant was filtered (0.22□μm). Serial dilutions of 5 μL of the filtrate were spotted on a *P. aeruginosa* lawn, and after overnight incubation at 37°C, the clearest plaques were picked, and their titer was determined^10^.

Five isolation streaks of each phage on its target bacterium lawn were performed to ensure that each phage was pure. Following this, phages were grown in liquid cultures with logarithmic-phase *P. aeruginosa* to a concentration of 10^8^ PFU/mL. Phages were stored at 4°C, and stocks of both phage and target bacterium were kept in 25% glycerol at -80°C.^8^

### Plaque forming unit (PFU) count

For the plaque-forming assay, phages were diluted 10-fold in LB broth and spotted over a PA bacterial lawn grown on soft 0.6% agar, and incubated overnight at 37°C. The observed plaques were counted, and the concentration of PFU was calculated^11^.

### Growth Curve Analysis

Lytic activity was evaluated using logarithmic bacteria (10^7^ CFU/ml) with the each phage at a concentration of 10^8^ PFU/ml and measuring turbidity using a plate reader (Synergy; BioTek, Winooski, VT) at OD600nm. The measurements were taken at 37°C with 5-second shaking intervals every 20 minutes for a duration of 24 hours^11^.

### Transmission Electron Microscopy (TEM)

Phage structure was visualized by Transmission Electron Microscopy (TEM) as described before.^10^ Briefly, 1 ml of 10^8^ PFU was centrifuged at 20,000 g (centrifuge 5430 R, rotor FA-45-24-11HS; Eppendorf, Hamburg, Germany) for 2h at room temperature. The pellet was resuspended in 200 μl of 5 mM MgSO_4_, spotted on a 2% uranyl acetate carbon-coated copper grid and incubated for 1 min. Phage visualization was carried out with a TEM 1400 plus (Joel, Tokyo, Japan) and images were captured with a charge-coupled device camera (Gatan Orius 600).

### Phage DNA purification and Sequencing

DNA from the phages was purified using a phage DNA isolation kit (Norgen Biotek). Library preparation and sequencing were conducted using the Illumina NextSeq 500 platform as described^10^ with single reads of 150bp with coverage >200.

### Genomic bioinformatic analysis

The reads were processed, and the genomes were assembled using Geneious Prime v2024.0.3 and its associated plugins. Briefly, the reads were trimmed using the BBDuk plugin and de-novo assembled with the SPAdes plugins^10^ and Geneious assembler. The assembled genomes were annotated using Pharokka and Phold.^12^ Topology was determined by both the presence of repeats at the beginning and end of the sequences, indicating their potential connection, and the repeated sequencing of a small group of reads. In a circular genome, the beginning and end of the sequence vary, while linear phages show a constant result. Virulence factors were identified using the VFDB database.^13^ Lysogenicity was determined by the presence of a C-like repressor gene and the phage’s similarity to bacterial chromosomes in BLAST analysis. Taxonomy was determined by the closest related phage found through BLAST analysis. ORF Genome map presentation was done using https://proksee.ca/^14^. VirClust was applied to cluster bacteriophages based on their protein profiles^15^. The nucleotide similarity of the phages was determined using Genious Prime, and the Orthologous Average Nucleotide Identity Tool (OAT; v0.93.1)^16^, which measures the overall similarity between two genome sequences.

## Results

Six phages were isolated from different sample from a sewage facility near Jerusalem, for a study of phage efficacy: PAShipCat1, PAKlein3, PACT201, PADD, PAB1, and PAB2. Their lytic activity, morphology, and genomes were assessed.

### Morphological Features

All six phages are tailed and exhibit distinct morphological characteristics. PAShipCat1, PAKlein3, and PACT201 display a Siphoviridae morphology with long, non-contractile tails. In contrast, PADD, PAB1, and PAB2 exhibit Myoviridae morphology, characterized by long, contractile tails (Figure 1 A-F)^17^.

**Figure 1:**
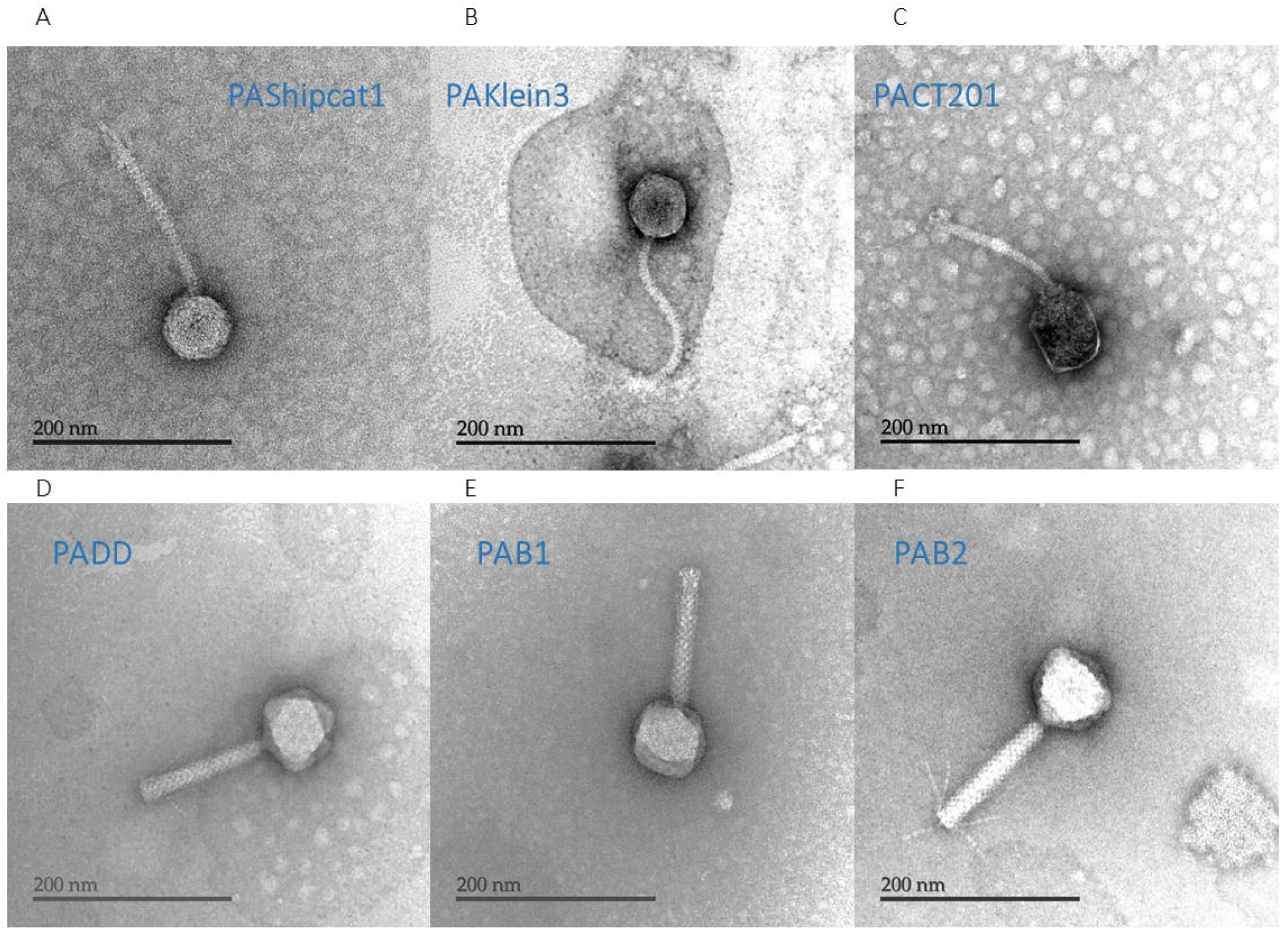
Morphological characterization of *Pseudomonas aeruginosa* phages. Transmission Electron Microscopy (TEM) images of the isolated *Pseudomonas aeruginosa* phages from the Israeli Phage Bank. (A) PAShipCat1, (B) PAKlein3, and (C) PACT201, exhibiting the typical Siphoviridae morphology with long, non-contractile tails. (D) PADD, (E) PAB1, and (F) PAB2, displaying the Myoviridae morphology characterized by long, contractile tails. Scale bars represent 200 nm.

### Lytic Activity

All six phages demonstrated lytic activity when tested in vitro using a plate reader on their host bacteria (Supplemental Figure 1A-F). PAB1 was the most effective phage, while PAShipCat1 was the least effective. Although PAB1 and PADD are genetically close, their lytic activity is non similar, PAB1 regrowth time is 1 hour after PADD.

### Genomic Features

PAKlein3, PAB1, and PAB2 have linear genomes while PAShipCat1, PADD, and PACT201 have genomes may be in a circular or linear topology (Table 1). BLAST analysis indicates that PAShipCat1 and PACT201 may have lysogenic potential due to their similarity to bacterial chromosomes. None of the six phages contain known virulence or antibiotic resistance genes, suggesting their potential suitability for phage therapy, particularly for the lytic phages PAKlein3, PADD, PAB1, and PAB2.

The genome sizes of these phages range from 45 kb (PAB2) to 67 kb (PADD) (Table 1, Figure 2 A-F). The GC content is highest among PA01-targeting phages, with PAKlein3 having 63.4%, compared to PA14-targeting phages, which have GC contents ranging from 55.6% (PAB1) to 59.4% (PAShipCat1) (Table 1, Figure 2 A-F). In contrast, PA14 and PA01 both have high GC contents of 66.1%^18^ and 67%^19^, respectively.

**Figure 2:**
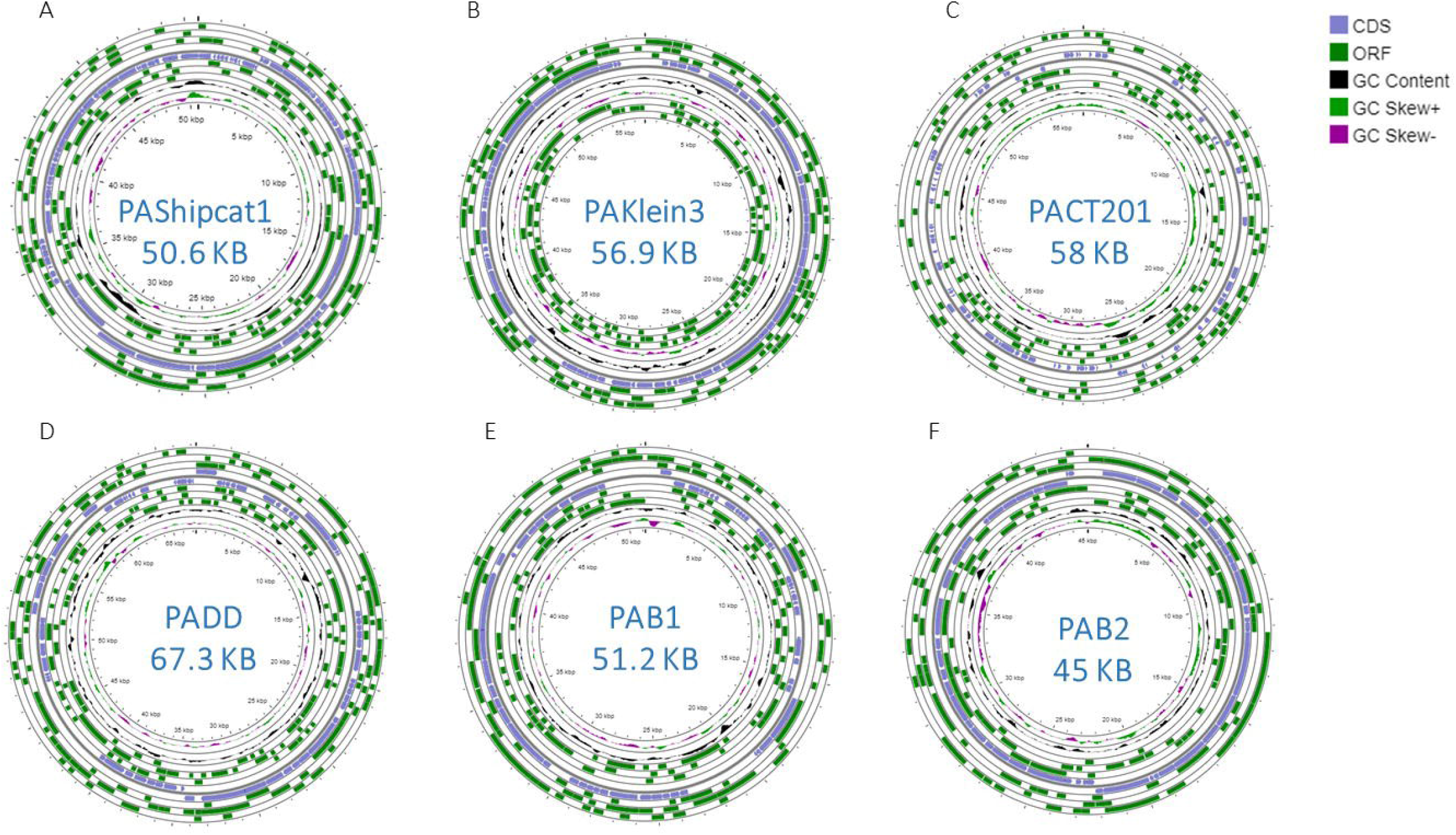
Genome maps of six phages, each showing their ORFs. (A) PAShipCat1, (B) PAKlein3, and (C) PACT201. (D) PADD, (E) PAB1, and (F) PAB2. The genomic maps were prepared using the CGView Server. The outer lane displays genes on the forward strand, while the next lane shows genes on the reverse strand, positive and negative GC Skew are presented.

While most phages have comprehensive genome annotations, PACT201 stands out due to its unique characteristics. Its Open Reading Frames (ORFs) are shorter than those of other phages, resulting in less detailed annotations. PACT201 is highly strain-specific and warrants further investigation (Figure 2).

### Phylogenetic analysis

Phages are grouped into three clusters based on genomic and protein similarities (Figure 3 A-C). The cluster of PACT201 and PAShipCat1 share potential lysogenic capabilities, and both are part of the Caudoviricetes taxonomy, the tailed phages taxonomy^20^. Another cluster of PAKlein3 and PAB2 showed close evolutionary relationships with linear genomes and no lysogenic markers, but they are of different taxonomy (Figure 3 A-C, Table 1). Their grouping suggests a purely lytic lifecycle, aligning with their therapeutic potential The highest similarity is observed between PADD and PAB1. Their genomic distance is 98.97%, measured by Average Nucleotide Identity (ANI), and both share a taxonomy within the Pbunavirus family (Figure 3 A-C, Table 1). On the protein level, their similarity is 85%. These phages have been isolated from different environmental samples, and therefor are considered different phages. The main genomic difference between these two phages is the presence of a glutamine amidotransferase gene on PADD which is not present in PAB1, and the presence of two hypothetical proteins on PAB1 which are not present on PADD (Figure 3 A-C).

**Figure 3:**
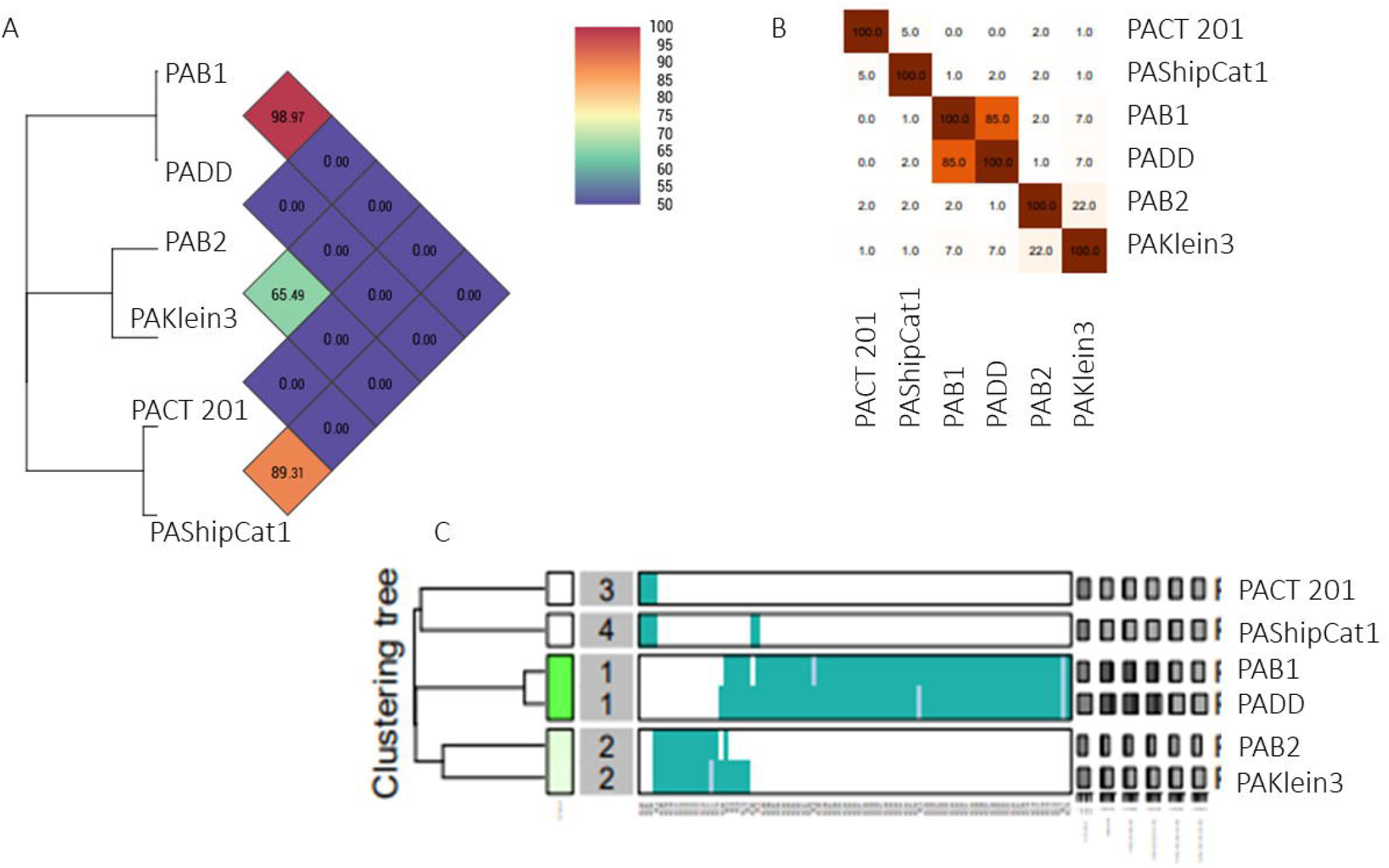
Comparative genomic analysis of bacteriophages. (A) Average Nucleotide Identity (ANI) among bacteriophage genomes. (B) Distance heatmap illustrating the relationships among bacteriophage genomes. (C) Clustering of viral bacteriophage proteins into two hierarchical levels. Proteins are initially grouped into protein clusters (PCs) based on their reciprocal BLASTP similarities. These PCs are subsequently organized into protein superclusters based on their hidden Markov model (HMM) similarities.

## Discussion

The genomic characterization of the six *Pseudomonas aeruginosa*-targeting bacteriophages—PAShipCat1, PAKlein3, PACT201, PADD, PAB1, and PAB2— provides significant insights into their potential applications in phage therapy.

The variations in genome topology and size, as well as GC content, reflect the diverse nature of these phages. Specifically, PAShipCat1 and PACT201 exhibit circular genomes and show potential lysogenicity due to their similarity to bacterial chromosomes in BLAST analysis.^21^ In contrast, the linear genomes of PAKlein3, PADD, PAB1, and PAB2, along with their absence of lysogenic markers, suggest a purely lytic lifecycle, which is advantageous for therapeutic applications.

The analysis of phage genomes using plain nucleotide similarity and Average Nucleotide Identity (ANI) offers distinct insights into genetic relatedness^16^. While two phages (PAB1 and PADD) are highly similar genetically, they show different lytic activity and are isolated from different samples and therefor considered different.

The reduced activity of PADD in comparison to PAB1 might be due to the presence of the glutamine amidotransferase (GATase) gene, required for correct decoding of glutamine codons during□translation^22^. The expression of GATase might impose an additional metabolic burden on the host cell, making it more challenging for the host to meet the demands of phage replication. This genetic load can result in increased cellular resource consumption and energy expenditure required for the replication and transcription of the additional genetic material. Consequently, the added metabolic demands and the strain on the host’s metabolic capacity could significantly impact the overall replication efficiency of the phage carrying the GATase gene^23^.

The morphological differences observed through Transmission Electron Microscopy (TEM) further distinguish these phages. The *Siphoviridae* morphology of PAShipCat1, PAKlein3, and PACT201, characterized by long, non-contractile tails, contrasts with the *Myoviridae* morphology of PADD, PAB1, and PAB2, which possess long, contractile tails^17^. These structural variations may influence their interaction with bacterial hosts and their efficacy in different environmental conditions.

The observation that the least effective phage, PAShipCat1, also exhibits lysogenic potential is consistent with the expectation that lysogenic phages may have reduced lytic activity compared to strictly lytic phages^24^. Lysogenic phages integrate into the host genome and establish a dormant state, which can limit their ability to cause cell lysis. This reduced lytic efficiency is a known trait of lysogenic phages, supporting the idea that their primary function may be related to latency rather than active infection. Understanding this relationship helps to clarify the functional roles of different phages and their potential applications in phage therapy^11^.

Our findings align with previous studies that highlight the potential of bacteriophages as a viable alternative to traditional antibiotics, especially against antibiotic-resistant strains of *P. aeruginosa*.^25,26^ The detailed genomic and morphological characterization provided in this study lays the groundwork for future research on the therapeutic efficacy and safety of these phages in clinical settings.

In conclusion, the six *Pseudomonas aeruginosa*-targeting bacteriophages described in this study exhibit promising features for phage therapy, including the absence of harmful genes, diverse morphologies, and adaptability to different bacterial strains. Further *in vivo* studies and clinical trials will be essential to validate their efficacy and safety for treating *P. aeruginosa* infections.

## Data Availability

The annotated genome sequences of the phages are available in GenBank under the accession numbers provided in the text (https://www.ncbi.nlm.nih.gov/genbank/).

## Acknowledgments

We thank the Israeli Phage Bank for providing the phage samples and the Core Research Facility team for their assistance with deep sequencing and TEM microscopy.

## Funding

The Milgrom Family Fund Grant #3015005877(AR, RH)

The Israel Science Foundation IPMP Grant #ISF1349/20 (RH)

The Rosetrees Trust Grant A2232 (RH).

The Foulks Foundation scholarship for MD-PhD students (AR).

## Author contributions

All authors made significant contributions to the preparation of this manuscript. AR, and RH conceptualized and drafted the manuscript, overseeing the funding acquisition process. SCG have edited the manuscript. OY, JB and SAO, were instrumental in executing various methodologies. Supervision throughout the project was provided by RH.

## Author Disclosure Statement

No competing financial interests exist.

During the preparation of this work, the authors used OpenAI-ChatGPT4 (https://chatgpt.com) Grammarly (https://app.grammarly.com) to correct language and grammar. After using this tool/service, the authors reviewed and edited the content as needed and took full responsibility for the content of the published article.

## Notes

### Competing Interest Statement

The authors have declared no competing interest.

## References

1. Gellatly SL, Hancock REW. Pseudomonas aeruginosa: new insights into pathogenesis and host defenses. Pathog Dis. 2013;67(3):159–73.

2. Nir-Paz R, Gelman D, Khouri A, et al. Successful treatment of antibiotic-resistant, poly-microbial bone infection with bacteriophages and antibiotics combination. Clinical Infectious Diseases. 2019;69(11):2015–8.

3. Onallah H, Hazan R, Nir-Paz R, et al. Refractory Pseudomonas aeruginosa infections treated with phage PASA16: A compassionate use case series. Med. 2023;4(9):600–11.

4. Salmond GPC, Fineran PC. A century of the phage: past, present and future. Nat Rev Microbiol. 2015;13(12):777–86.

5. Chegini Z, Khoshbayan A, Taati Moghadam M, et al. Bacteriophage therapy against Pseudomonas aeruginosa biofilms: A review. Ann Clin Microbiol Antimicrob. 2020;19:1–17.

6. Phee A, Bondy-Denomy J, Kishen A, et al. Efficacy of bacteriophage treatment on Pseudomonas aeruginosa biofilms. J Endod. 2013;39(3):364–9.

7. Braunstein R, Hubanic G, Yerushalmy O, et al. Successful phage-antibiotic therapy of P. aeruginosa implant-associated infection in a Siamese cat. Veterinary Quarterly. 2024;44(1):1– 9.

8. Yerushalmy O, Khalifa L, Gold N, et al. The Israeli phage bank (IPB). Antibiotics. 2020;9(5):269.

9. Swetha K, Stefan KL, A. Og. Differential Surface Competition and Biofilm Invasion Strategies of Pseudomonas aeruginosa PA14 and PAO1. J Bacteriol. 2022 Mar 19;203(22):e00265-21. Available from: 10.1128/JB.00265-21

10. Rimon A, Rakov C, Lerer V, et al. Topical phage therapy in a mouse model of Cutibacterium acnes-induced acne-like lesions. Nat Commun. 2023;14(1):1005.

11. Yerushalmy O, Braunstein R, Alkalay-Oren S, et al. Towards Standardization of Phage Susceptibility Testing: The Israeli Phage Therapy Center “Clinical Phage Microbiology”—A Pipeline Proposal. Clinical Infectious Diseases. 2023;77(Supplement_5):S337–51.

12. Bouras G, Nepal R, Houtak G, et al. Pharokka: a fast scalable bacteriophage annotation tool. Bioinformatics. 2023;39(1):btac776.

13. Altschul SF, Madden TL, Schäffer AA, et al. Gapped BLAST and PSI-BLAST: a new generation of protein database search programs. Nucleic Acids Res. 1997;25(17):3389–402.

14. Grant JR, Enns E, Marinier E, et al. Proksee: in-depth characterization and visualization of bacterial genomes. Nucleic Acids Res. 2023;51(W1):W484–92.

15. Moraru C, Varsani A, Kropinski AM. VIRIDIC—A novel tool to calculate the intergenomic similarities of prokaryote-infecting viruses. Viruses. 2020;12(11):1268.

16. Lee I, Ouk Kim Y, Park SC, et al. OrthoANI: an improved algorithm and software for calculating average nucleotide identity. Int J Syst Evol Microbiol. 2016;66(2):1100–3.

17. Linares R, Arnaud CA, Degroux S, et al. Structure, function and assembly of the long, flexible tail of siphophages. Curr Opin Virol. 2020;45:34–42.

18. Lee DG, Urbach JM, Wu G, et al. Genomic analysis reveals that Pseudomonas aeruginosa virulence is combinatorial. Genome Biol. 2006;7:1–14.

19. Grocock RJ, Sharp PM. Synonymous codon usage in Pseudomonas aeruginosa PA01. Gene. 2002;289(1–2):131–9.

20. Turner D, Kropinski AM, Adriaenssens EM. A roadmap for genome-based phage taxonomy. Viruses. 2021;13(3):506.

21. Feiner R, Argov T, Rabinovich L, et al. A new perspective on lysogeny: prophages as active regulatory switches of bacteria. Nat Rev Microbiol. 2015;13(10):641–50.

22. Curnow AW, Hong K won, Yuan R, et al. Glu-tRNAGln amidotransferase: a novel heterotrimeric enzyme required for correct decoding of glutamine codons during translation. Proceedings of the National Academy of Sciences. 1997;94(22):11819–26.

23. Howard-Varona C, Lindback MM, Bastien GE, et al. Phage-specific metabolic reprogramming of virocells. ISME J. 2020;14(4):881–95.

24. Howard-Varona C, Hargreaves KR, Abedon ST, et al. Lysogeny in nature: mechanisms, impact and ecology of temperate phages. ISME J. 2017;11(7):1511–20.

25. Altamirano FLG, Barr JJ. Phage therapy in the postantibiotic era. Clin Microbiol Rev. 2019;32(2).

26. Strathdee SA, Hatfull GF, Mutalik VK, et al. Phage therapy: From biological mechanisms to future directions. Cell. 2023 Jan 5;186(1):17–31. Available from: 10.1016/j.cell.2022.11.017

27. Grace A, Sahu R, Owen DR, et al. Pseudomonas aeruginosa reference strains PAO1 and PA14: A genomic, phenotypic, and therapeutic review. Front Microbiol. 2022;13:1023523.

