## Supplemental data for "Complete Genome Sequence of Six *Pseudomonas aeruginosa* Bacteriophages aim for research and treatments"

**Supplementary material Rimon et al.**

**Figure legend.**

**Supplemental Figure S1.** Growth curves of all phages on planktonic bacteria (A-E).

Optical density (OD 600nm) of planktonic *P. aeruginosa* treated with six different phages at a concentration of 10^8^ PFU/ml, corresponding to MOI of 10 (A) PAShipCat1, (B) PAKlein3, and (C) PACT201. (D) PADD, (E) PAB1, and (F) PAB2. The results are the average of triplicates, presented as mean ± standard deviation.

**Supplemental Figure S1.**

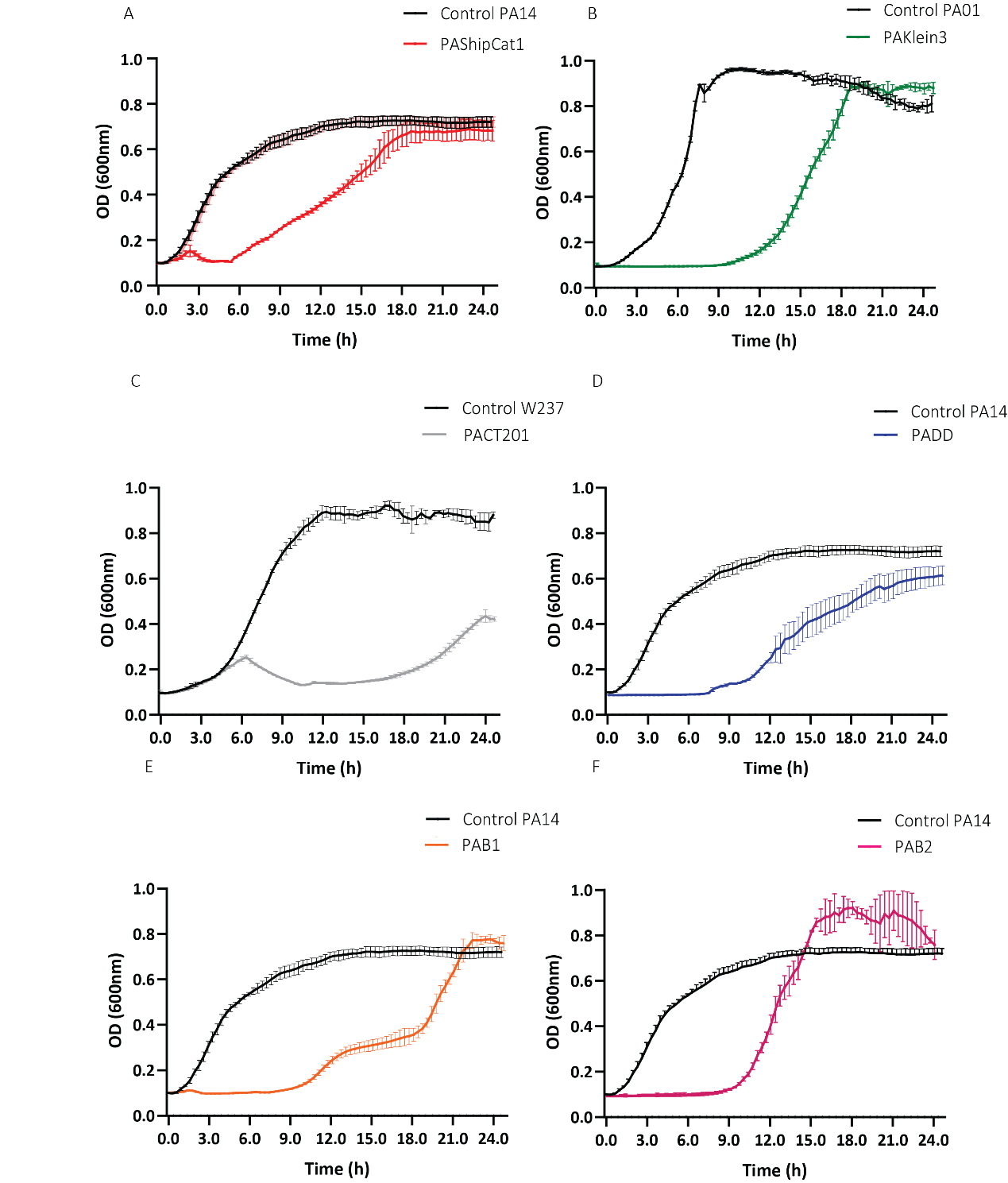
